# Membrane molecule bouncer enables follicle fertilization in a viviparous teleost: *Poecilia reticulata* (guppy)

**DOI:** 10.1101/2023.08.02.551741

**Authors:** Junki Yoshida, Yuki Tajika, Kazuko Uchida, Makoto Kuwahara, Kaori Sano, Takayuki Suzuki, Eiichi Hondo, Atsuo Iida

## Abstract

Fertilization is a fundamental mechanism of sexual reproduction. Generally, oocytes are ovulated from the ovarian follicles and contact the sperm outside the ovarian medulla. Unlike this, follicle fertilization which means the egg contact with sperm in the ovarian medulla without ovulation is known in the viviparous teleost species belonging to the Poeciliidae. In this study, we focused on a viviparous teleost species, *Poecilia reticulata* (guppy). Our sperm tracking assay indicated that the sperm reached the immature oocytes with a germinal vesicle, and the fertilized immature oocytes were presumed to contribute to littermates. The binding between immature oocytes and sperm is a specific trait in the guppy, which was not observed in *Danio rerio* (zebrafish) or *Oryzias latipes* (medaka). The loss- and gain-of-function assays indicated that bouncer plays a critical role in immature oocyte-to-sperm binding. This fertilization trait in immature oocytes may provide certain advantages for females with respect to nutrition or other gestation costs. Our findings shed light on the unique reproductive strategies of guppy and contribute to our understanding of the diverse reproductive mechanisms in vertebrates.

**Summary statement:** Unlike general vertebrates, guppy’s oocyte fertilizes with sperm in the ovarian medulla at the immature stages. The distinctive trait depends on the Ly6/uPAR protein bouncer.

## Introduction

Fertilization is a fundamental mechanism for generating offspring in multicellular organisms [1]. Fertilization consists of multiple steps, including the approach of the gametes, binding of sperm to the oocyte, entry of the sperm nuclei into the oocyte, fusion of the sperm and oocyte nuclei, and activation of the fertilized egg [2,3]. Fertilization generally occurs outside the ovarian medulla after ovulation in both viviparous and oviparous animals [4]. In most mammals and fishes, ovulated secondary oocytes possess fertilization capacity and develop into mature eggs via a second polar body release after fertilization [5,6].

However, the prevalence of fertile oocyte stages across animals remains unclear. *Poecilia reticulata* (guppy) is a viviparous teleost species belonging to the family Poeciliidae (order Cyprinodontiformes) and is prevalent in freshwater fields in South and Middle America [7]. The male possesses a modified anal fin structure as external genitalia called the gonopodium and uses it as an intromittent organ for internal fertilization [8,9]. In the ovary, oocytes are kept in the follicle until the later stage of embryogenesis, and narrow ducts connecting the ovarian lumen to the follicles develop. The sperm ejected from the male into the female ovary approaches the oocytes through the ducts and fertilizes them in the follicle [10,11].

Sperm–oocyte binding is not only the first step of fertilization but also one of the gatekeepers to avoid crossbreeding between different species [12,13]. In invertebrates, including sea urchins and abalones, zona pellucida proteins present on egg surfaces play important roles in prevention of crossbreeding [14]. Juno, a glycosylphosphatidylinositol (GPI) anchor protein present on egg surfaces, and Izumo1, an immunoglobulin transmembrane protein present on the sperm head, are coupled and are necessary for fertilization in mammals [15]. In teleosts, bouncer, a GPI anchor protein expressed in eggs, and sperm acrosome associated-6 (spaca6), a transmembrane protein expressed in sperms, are known to be responsible for sperm–oocyte binding; however, whether these molecules couple with each other remains unknown [16,17]. The amino acid sequences of bouncers are diverse among teleosts, and bouncers are considered to be responsible for regulating species-specific sperm–oocyte binding in teleosts [16]. On the other hand, a recent study indicated the molecular signatures that defined species specificity and discussed the cross-fertility between the teleost species [18].

In this study, we focused on the ovarian structure and fertility timing of a viviparous teleost species, guppy, in which oocytes are fertilized by sperms in the ovarian medulla without ovulation. We hypothesized that oocyte fertility is restricted by duct development and not by oocyte stage or ovulation in this species. To solve the question, we investigated sperm– oocyte binding, which is the first step in fertilization. We aimed to evaluate the physiological requirement for follicle fertilization, and its responsible factor focusing on bouncer/spaca4l genes in guppy.

## Results

### Sperms can fertilize immature oocytes in guppy

Histological analysis indicated that no ovulated oocytes were present in guppy ovaries.

Narrow ducts connecting the ovarian lumen to the oocytes were present. The ducts were connected to not only matured oocytes post vitellogenesis but also immature oocytes of 50–100 µm diameters (Figure 1A and B, Movie S1). The ducts, made of epithelial layer cells, continued into the ovarian luminal epithelium, and the proximal end opened into the granule membrane surrounding the oocytes (Figure 1C). To validate whether the sperm reaches immature oocytes, tracer analysis was performed by injecting nucleus-labeled sperm into the ovarian lumen from the genital pore (Figure 1D). The spermatozoa were stained with Hoechst while maintaining their swimming capacity (Figure 1E). Hoechst-positive signals that indicate the injected spermatozoa were observed on the surface of the immature oocytes (Figure 1F). Furthermore, electron microscopy revealed the invasion of sperm into the ducts (Figure 1G). To estimate the fertility of immature oocytes at the pre- vitellogenesis stage, we conducted artificial fertilization, in which the number of mature oocytes in non-pregnant females was compared to that of the littermates obtained as a result of natural delivery after sperm injection (Figure 1H). The average number of littermates per injection was significantly higher than that of mature oocytes in single ovaries of non-pregnant females (Figure 1I). Furthermore, two of the five females became pregnant again without additional sperm injections or mating after primary delivery (Table S1, 2).

**Figure 1.**
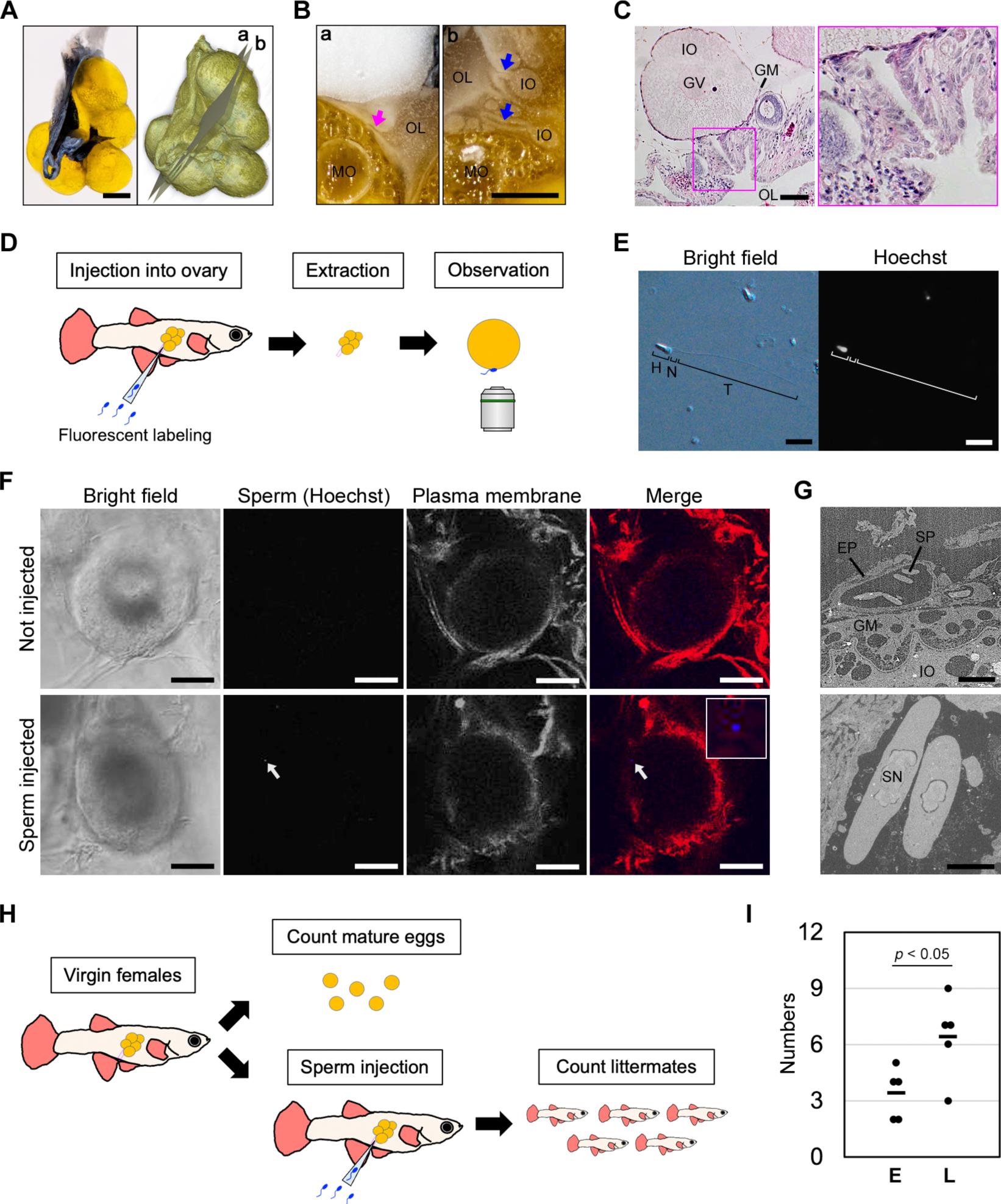
Sperm binding and fertilization activity in immature oocytes. **A**. A macroscopic image of *P. reticulata* ovary. A volume rendered image were made from 1687 serial block-face images. Scale bar, 1 mm. **B**. In the block-faces at the planes a and b, ducts between the ovarian lumen and mature or immature oocytes (magenta and blue arrows, respectively) are seen. IO: immature oocyte, MO: mature oocyte, OL: ovarian lumen. Scale bar, 500 µm. **C**. HE-stained section for the duct structure through the immature oocyte. GM, granule membrane. GV, germinal vesicle. Scale bar, 50 µm. **D**. Schematic representation of our artificial fertilization procedure used to observe sperm binding with oocytes. **E**. Fluorescence images for sperms stained using Hoechst. The nucleus was visualized using a filter for blue fluorescence. H, head. N, neck, T, tail. Scale bar, 10 µm. **F**. Confocal microscopy used to detect the Hoechst- labeled sperm on the surface of an immature oocyte. The white arrow indicates the sperm nucleus. Scale bar, 50 µm. **G**. Electron microscopy for the artificially injected sperm in the cavity of the duct. EP, epithelial cells, GM, granule membrane. IO, immature oocyte. SN, sperm nucleus. SP, sperm. Scale bars, 10 µm (upper), 2 µm (lower). **H**. Schematic representation of our artificial fertilization procedure used to count the mature oocytes in the ovary and the littermate number of offspring. **I**. Comparison between the number of mature oocytes in the ovary and the delivered offspring among the littermate females. The littermate number is significantly higher than the matured ovary number, as indicated by the Student’s t-test. E, egg. L, littermate.

### Guppy sperms do not bind to oocytes heterogeneously

To validate whether binding between sperm and immature oocytes is a specific trait in the guppy, an *in vitro* sperm binding assay was performed using the oviparous species zebrafish *Danio rerio* (zebrafish) and *Oryzias latipes* (medaka) (Figures 2A). Guppy oocytes were observed to bind to sperms under *ex vivo* conditions. In contrast, the immature oocytes of oviparous fish exhibited no binding capacity to the sperm isolated from mature males of each species (Figure 2B). Next, to validate whether guppy oocytes or sperms possess the nonspecific binding capacity to any other cell, a swapping assay in which guppy oocytes were co-incubated with medaka sperm or vice versa was performed (Figure 2A). The guppy gametes did not exhibit interspecies binding in the sperm binding assay (Figure 2B). Thus, our study revealed that sperm-binding capacity in immature oocytes was a unique characteristic in guppy.

**Figure 2.**
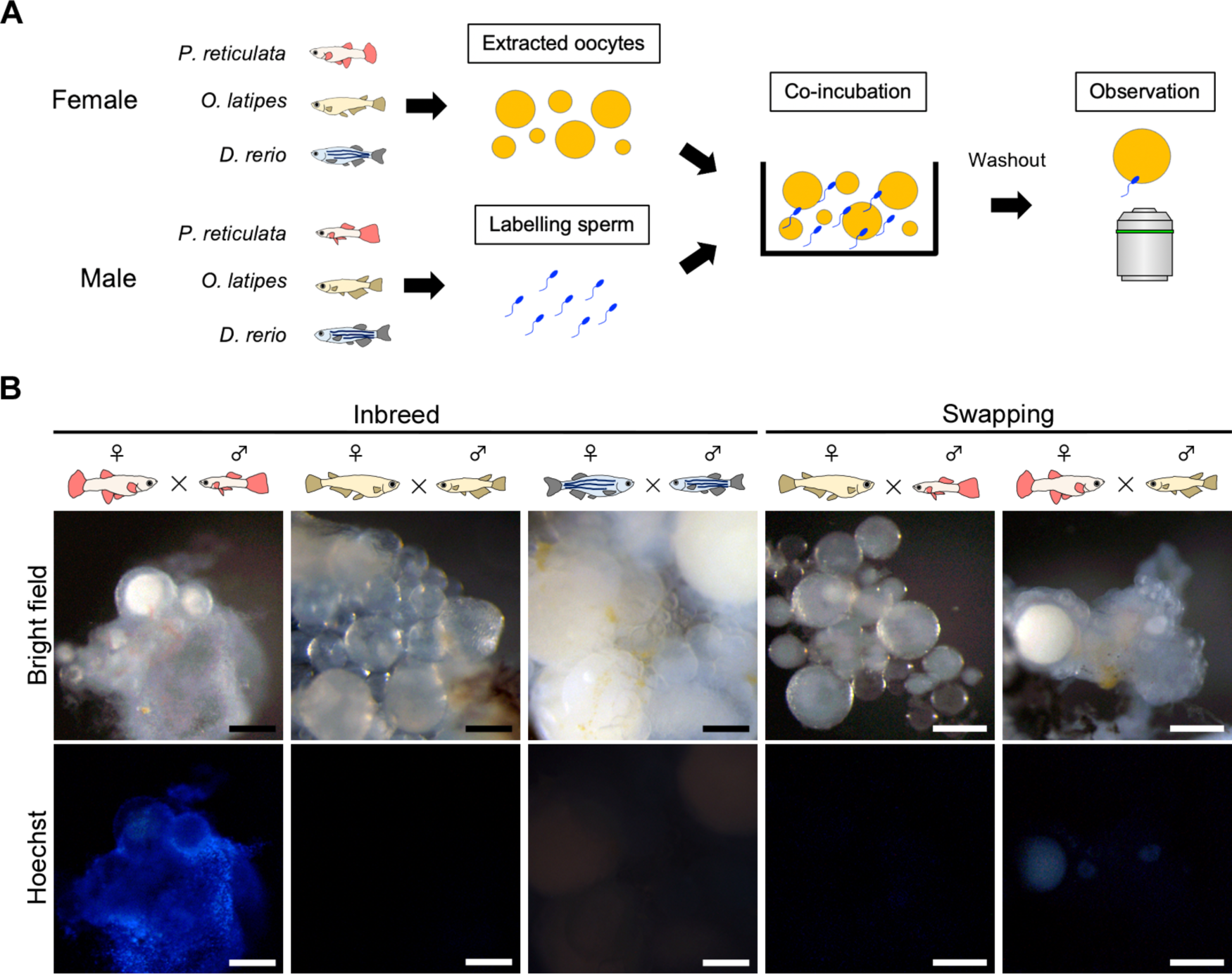
Species specificity of sperm-binding capacity to immature oocytes. **A**. Schematic representation of the *in vitro* sperm binding assay. Sperms and oocytes were isolated from guppy, medaka, zebrafish respectively. The sperms were fluorescent-labeled and co-incubated with the oocytes. After the washing, signals for the oocyte-bound sperm were observed using fluorescent microscopy. **B**. Fluorescent microscopy for the ovaries after the insemination. Scale bar, 200 µm.

### Cloning and characterizing of guppy bouncer/spaca4l

To explore a factor responsible for binding between immature oocytes and sperm in the ovary of guppy, we focused on the bouncer, which is involved in egg–sperm binding in oviparous teleosts, including zebrafish. Based on the phylogenetic analysis, a GPI anchor protein-coding gene registered as *spaca4l* (NCBI: XP_008410152.1) was used as a candidate that is closely related to the zebrafish bouncer (Figure 3A). The *in situ* hybridization indicated that the guppy *bouncer* was expressed in oocytes of ovarian follicles. Particularly, the expression was higher in the immature oocytes with a diameter in the range of 50–100 µm with a germinal vesicle than in the mature oocytes post vitellogenesis (Figure 3B). Phylogenetic analysis indicated that the guppy bouncer is closely related to the zebrafish and medaka bouncers that were reported to function in the oocyte–sperm binding capacity in a previous study [16] (Figure 3C). The amino acid sequences of the bouncers were diverse among species; however, the cysteine (C) loci that form disulfide bonds were highly conserved. Glycosylation motifs in the finger structure observed in the zebrafish or medaka bouncers were not included in the guppy bouncer (Figure 3D).

**Figure 3.**
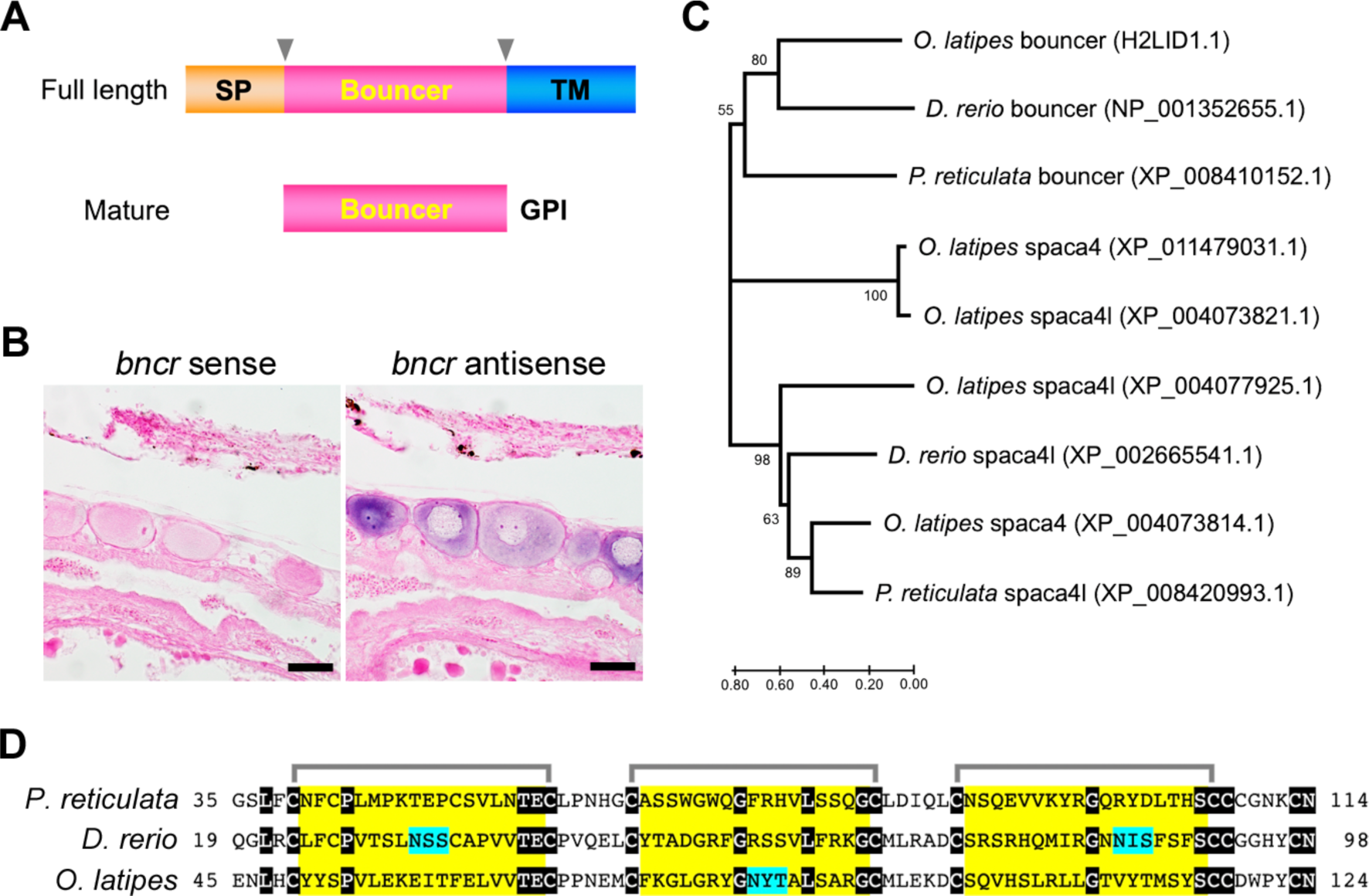
Identification of bouncer protein in *P. reticulata*. **A**. Typical structure of the teleost bouncer protein. The signal peptide (SP) and transmembrane (TM) are cleaved off from the full-length protein. The functional bouncer protein is localized on the cell surface with the help of a glycosylphosphatidylinositol (GPI) anchor. **B**. The distribution of bouncer transcripts by *in situ* hybridization. The strong signals were observed in the immature oocytes with germinal vesicles. Scale bar, 50 µm. **C**. A phylogenetic tree of spaca4l and bouncer protein of *P. reticulata*, *O. latipes*, and *D. rerio*. **D**. Alignment for the amino acid sequences of bouncer protein in *P. reticulata*, *O. latipes*, and *D. rerio*. The black background residues indicate conserved amino acids among the three species. The brackets indicate that cysteines consist of disulfide bonds. The yellow background residues indicate finger structure consists of the disulfide bonds that are the interface of receptor association. The light blue background triplicate residues indicate the predicted motifs for N-glycosylation (NXS/T. X is except proline.).

### Bouncer regulates oocyte–sperm binding in guppy

As a loss-of-function analysis of the bouncer for oocyte–sperm binding, a dominant negative assay using the soluble form of the bouncer protein (bouncer decoy) was performed. Guppy sperm preincubated with the bouncer decoy lacked the oocyte-binding capacity. In contrast, co-incubation with a negative control decoy derived from zebrafish bouncer did not interfere with the oocyte–sperm binding (Figure 4A). To perform the gain-of-function analysis, we prepared a transgenic zebrafish strain which expressed guppy bouncer specifically in the oocyte (Tg[*gdf9*:*prbncr-2a-gfp*]) (Figure 4B and C). We observed that transgenic zebrafish oocytes expressing guppy bouncer could bind to guppy sperm by the sperm binding assay. In contrast, oocytes of the negative control zebrafish strain (Tg[*gdf9*:*gfp*]) did not bind to the guppy sperm (Figure 4D).

**Figure 4.**
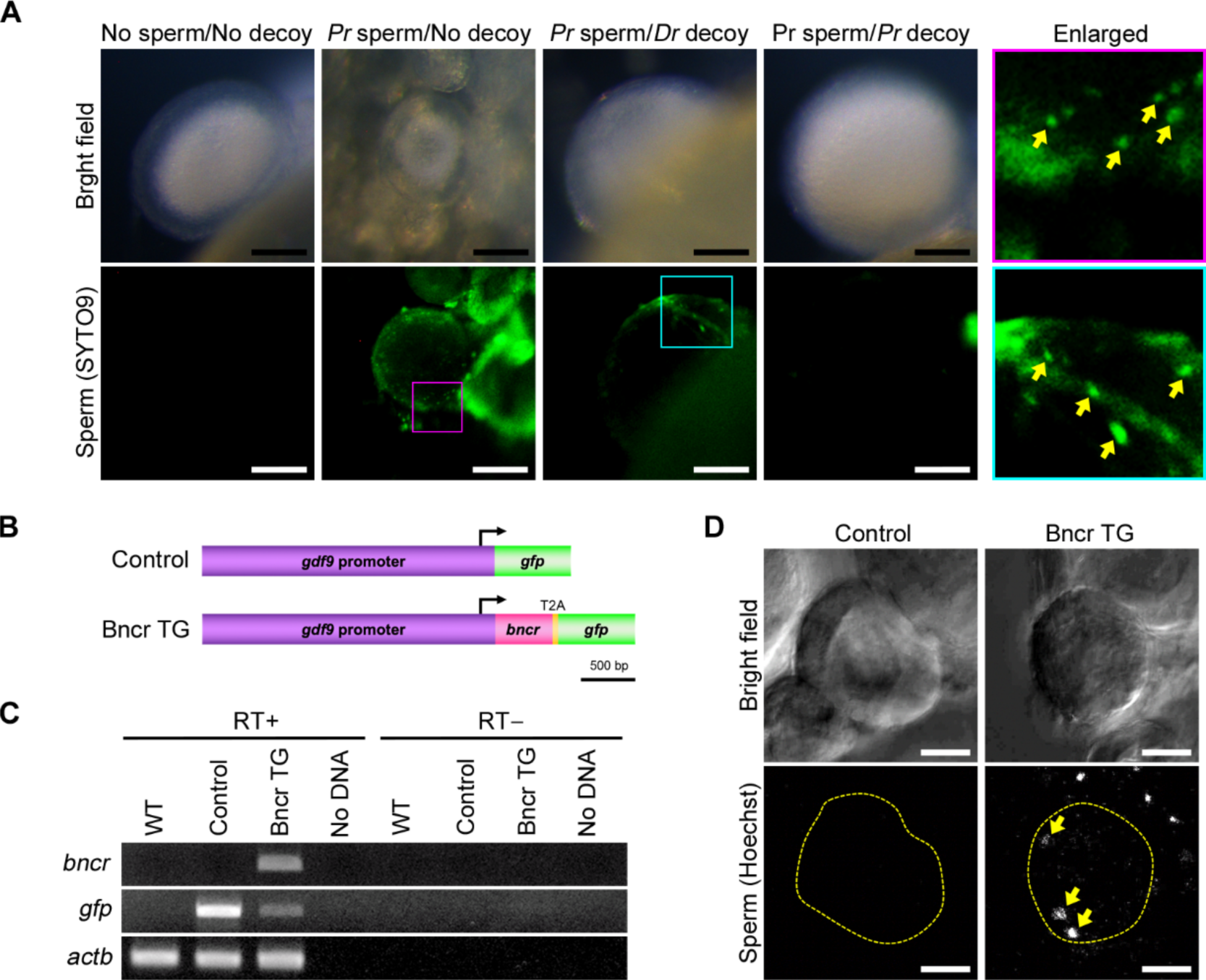
Functional analyzes of guppy bouncer. **A**. Microscopy images of the ovaries after the inbreeding sperm binding assay of guppy (*Pr*) with or without pretreatment of the recombinant decoy peptide obtained from guppy or zebrafish (*Dr*) bouncer sequences. **B**. DNA constructions for generation of transgenic zebrafish lines. The intact GFP (Control) or bouncer-T2A-GFP coding sequence is driven by the zebrafish *gdf9* promoter. **C**. RT-PCR used to detect the expression of the transgenes in the transgenic zebrafish lines. The *actb* was used as the internal control. The transgenes (*gfp* and *bncr*) were detected in the RT+ condition of the transgenic lines. **D**. In vitro sperm binding assay. The Hoechst-labeled guppy sperms on immature oocytes of the transgenic zebrafish strains were observed by confocal microscopy. The dotty lines and arrows indicate the oocyte contour and sperm nuclei, respectively. Scale bars, 20 µm.

## Discussion

In this study, we demonstrated the sperm-binding capacity in immature oocytes prior to vitellogenesis in guppy. Our artificial fertilization assay also suggests that immature oocytes bound to sperm contribute to the next generation of offspring. Furthermore, immature oocyte-to-sperm binding was observed only in guppy and not in zebrafish or medaka. Thus, early fertility is a trait specific to guppy. As a candidate molecule responsible for these traits, we focused on guppy *bouncer*/*spaca4l*, which is predominantly expressed in the early oocyte stages. In the loss-of-function assay, immature oocyte-to-sperm binding was disrupted by pretreatment with a decoy form of the guppy bouncer prior to the sperm binding assay. For the gain-of-function assay, we generated a transgenic zebrafish strain expressing guppy bouncer in immature oocytes and demonstrated that zebrafish oocytes expressing guppy bouncer possessed the capacity of binding to guppy sperm (Figure 5A). Our findings indicate that the bouncer is one of the factors responsible for immature oocyte-to-sperm binding in guppy.

**Figure 5.**
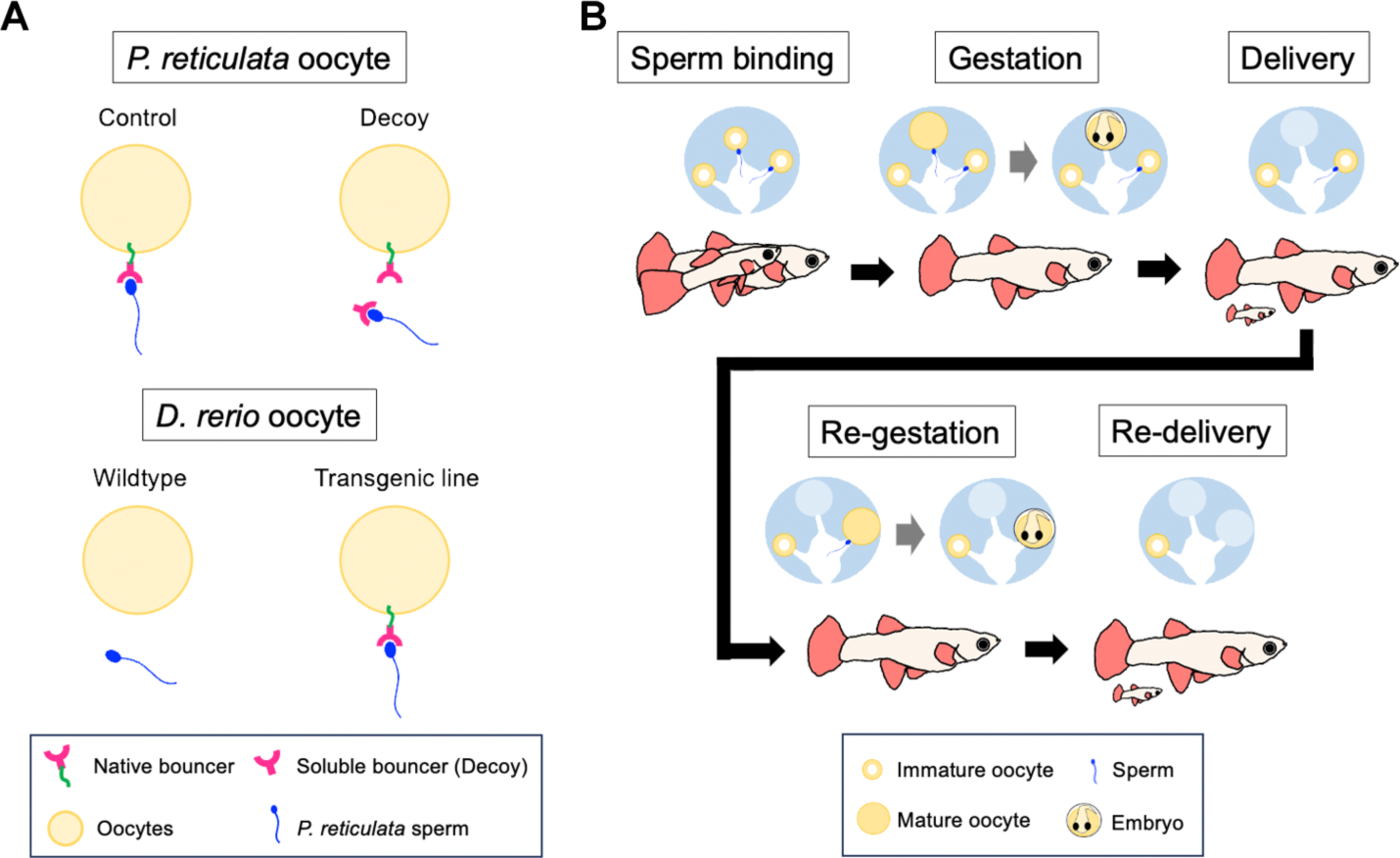
Model of the bouncer-mediated follicle fertilization process in the guppy ovary. **A**. A model of the bouncer-mediated sperm binding to immature oocytes based on our functional assay. In native conditions, prior to vitellogenesis, bouncer and the GPI-anchored protein expressed on immature oocytes contribute to sperm binding. The soluble bouncer protein works as a decoy to inhibit sperm binding. In zebrafish oocytes, bouncer proteins that possess an affinity to the guppy sperm are not provided. In the transgenic line, the guppy bouncer expressed in zebrafish oocytes as a transgene facilitates guppy sperm binding with the oocytes. **B**. A hypothesis for physiological roles of the sperm-binding capacity with immature oocytes in guppy.

Egg-to-sperm binding is a common phenomenon observed during the fertilization of most animals; however, the factors responsible for this binding vary among species. In *Strongylocentrotus purpuratus* (purple sea urchins), esr1 plays an important role in egg– sperm binding [19]. In *Mus musculus* (house mice), the coupling of Juno and Izumo1 is necessary for oocyte-to-sperm binding and subsequent fertilization steps [15,20]. In teleosts, bouncers in eggs and spaca6 in sperms, which are immunoglobulin proteins, play critical roles in egg-to-sperm binding in zebrafish [16,17]. However, the interaction between the bouncer and spaca6 remains unclear. In this study, our findings indicated that the unique fertilization style of sperm binding to intrafollicular oocytes prior to vitellogenesis in guppy is regulated by the spaca4l/bouncer protein expressed in immature oocytes. This suggests that intrafollicular fertilization, which is a trait specific to viviparous species belonging to the family Poeciliidae, occurs through modification of the temporal regulation of the bouncer gene expression. Additionally, we found that guppy immature oocytes possess no activity for binding to the medaka sperms and that the amino acid sequence of guppy bouncer was different from that of medaka or zebrafish. Therefore, it is suggested that the bouncer- dependent immature oocyte-to-sperm binding is a species-specific trait in guppy.

In almost all animals, complete maturations of eggs are not essential for sperm to fertilize eggs. In sea urchins, eggs are spawned and fertilized at the stage of female pronucleus formation after the second meiotic division. In *Caenorhabditis elegans*, sperms can fertilize eggs during germinal vesicle breakdown before reaching the miotic stage [21]. Even in vertebrates, including mammals, eggs are ovulated in the middle of the second meiotic division and can then be fertilized inside or outside the mother [22]. In these species, the immature oocytes are bind with sperm at the post-ovulation stages.

Fertilization of follicular oocytes had not been reported. Here we argue that follicular oocytes possess sperm-binding capacity and undergo fertilization in guppy and that the bouncer-defined sperm–oocyte binding provides not only species specificity but also variations of reproductive fashion including fertility timing in each species; however, the actual oogenetic stages that lead to fertility have remained unclear [10,11]. Furthermore, we only observed binding between oocytes and sperm; thus, the subsequent fertilization steps remain elusive within the previtellogenic stages.

The fertility in the previtellogenic stage of the host species has advantages. We postulate that the fertility of immature oocytes reduces the cost of oogenesis in females. Thus, females can maintain their eggs in the previtellogenic stage, and the eggs can then be developed after receiving the sperm. This method is different from the method in which all fertile eggs are kept as mature eggs with yolk. Female fish with this property possess advantages of reducing the cost of oogenic nutrients and predation risk during oogenesis and gestation. In viviparous species, the mother’s cost during gestation is a serious concern for the survival of female fish. In *Poecilia latipinna* (sailfin molly), resources available to the intraovarian embryo change depending on the nutritional conditions of the mother. [23]. In some viviparous teleost species belonging to the family Poeciliidae, superfetation reduces the negative effects of gestation, such as body shape changes and decreases in swimming ability [24]. We propose that fertility during the previtellogenic stage is a novel mechanism that reduces the costs of female reproduction. Additionally, sperm storage is another distinct trait in guppy to allow repetitive gestations resulting from a single mating [25]. This trait contributes not only to the genomic diversity of the offspring owing to the sperm received from multiple males in a single gestation but also to the reduction in the cost of exploring male fish as mating partners at the appropriate reproduction stages in depopulated environments. In our study, two female fish became pregnant twice as a result of a single sperm injection. We hypothesized that the immature oocyte-to-sperm binding capacity contributes to sperm storage (Figure 5B).

In summary, bouncer-mediated early fertility plays a role in the viviparous reproductive system in guppy by reducing female costs.

## Materials and Methods

### Animal experiment

This study was approved by the Ethics Review Board for Animal Experiments at Nagoya University (A210264-001). We euthanized a minimal number of live animals under anesthesia according to institutional guidelines. The live fish samples were euthanized using 0.01 % tricaine and on ice.

### Fish breeding

Guppy was purchased from Meito Suien Co., Ltd. (Nagoya, Japan). The Mic strain of zebrafish bred in our laboratory was used in this study. The HdrR strain of medaka was obtained from NBRP medaka (https://shigen.nig.ac.jp/medaka/). Adult fish were maintained in fresh water at 27 °C under a 14:10 L:D photoperiod cycle. Fertilized eggs to larval stage fish were maintained in a Petri dish containing 0.3% sea salt water.

### Block-face observation

The extracted ovaries were fixed in 4.0% paraformaldehyde/PBS (PFA/PBS) at 4 °C overnight. The fixed tissues were washed thrice with PBS. The ovaries were stained with 1% tannic acid/PBS for 2 h, washed with PBS, and embedded in white OCT compound. Image datasets of ovarian samples were obtained by correlative microscopy and block-face imaging [26, 27].

### Histology

The extracted tissues were fixed in 4.0% PFA/PBS at 4 °C overnight. Then, the fixed tissues were washed three times with PBS and then, dehydrated stepwise in 25, 50, 75, and 100% methanol. The tissues were degreased with xylene three times, then the solution was replaced with benzene, followed by replacement with 50% soft paraffin/benzene for 30 min. The samples were embedded in a paraffin block using soft paraffin (48 °C, 2 h) and then, hard paraffin (60 °C, 15 min, four times). The sample block was cut into 5 µm-thick slices and dried on gelatin-coated glass slides (overnight, 37 °C). The slides were deparaffinized with toluene; hydrated stepwise with 100, 95, 90, 85, and 70% ethanol; and then, stained with hematoxylin and eosin. The stained samples were dehydrated stepwise in 70, 80, 90, 95, and 100 % ethanol. After permeabilization with xylene, the specimens were embedded in Entellan™ New (Sigma-Aldrich, 107961) and observed under the BX53 upright microscope (Olympus, Tokyo, Japan).

### Electron microscopy

The tissue samples were fixed with 2.5% glutaraldehyde and then, washed with PBS. The immature oocytes were collected and embedded in 1.0% agar. The samples were washed twice with PBS on ice and treated with 1% osmium tetroxide/PBS for 80 min. Then, the samples were dehydrated stepwise using 50, 70, 80, 90, 95, 99.5, and 100% ethanol. The samples were treated with propylene oxide (Kishida) and then, embedded in epoxy resin (Nissin EM Quetol-812). The blocks were polymerized at 60 °C for 48 h. Sections of 100 nm thickness were made (Leica EM UC7), and stained with UranyLess (Micro to Nano) and 3% lead citrate (Micro to Nano). The backscattered electron images were obtained using a field-emission scanning electron microscope SU9000 (Hitachi, Tokyo, Japan) at 1kV.

### Sperm injection and tracing

Testes extracted from mature males were homogenized, and the lysates were stained with Hoechst 33342(DOJINDO, 23491-52-3) for 10 min at room temperature (RT, 25–27 °C) and then, washed three times with PBS. The lysates, including spermatozoa, were diluted in the Leibovitz L-15 medium (L-15, Gibco 11415064) and injected into female ovaries through the genital pore. The injected females were placed back in the breeding tank and maintained freely for 16 h. The ovaries were extracted and stained with CellMask (Invitrogen, 2491421) for 10 min. The tissue samples were washed with PBS three times and then, fixed in 4% PFA/PBS at 4 °C for 30 min. Immature eggs were surgically isolated from the tissue samples and then, embedded in 1% low-melting-point agarose in a glass bottom dish. To detect the signals indicating sperm bound to the egg surface, the samples were observed using the FV3000 Confocal Microscope (Olympus).

### Artificial fertilization assay

The testes extracted from mature males were homogenized. Lysates, including spermatozoa, were diluted in L-15. Virgin (non-pregnant) sibling females were anesthetized in ice water, the ovaries obtained from some of the siblings were extracted, and the number of mature eggs with diameters greater than 1 mm was counted. Lysates were injected through the genital pores into the other siblings. The injected female was put back in the tank and singly bred until the delivery (28–68 days, mean ± se = 46.3 ±5.04).

### In vitro sperm binding assay

Extracted immature oocytes and testis homogenates treated with Hoechst were co-cultured in L-15 in the same Petri dish for 10 min and washed with PBS three times. The oocytes were fixed in 4% PFA/PBS for 30 min and observed using the Leica M205FA stereomicroscope (Leica Microsystems, Wetzlar, Germany) or FV3000 Confocal Microscope.

### Cloning of guppy bouncer

A BLAST search based on zebrafish *bouncer* (NM_001365726.1) sequence information identified three *spaca4l* genes in the guppy genome. We focused on *spaca4l* (XM_008411930.1) as a guppy bouncer. Total RNA was extracted from adult guppy ovaries using the RNeasy Mini kit (QIAGEN) and reverse-transcribed using SuperScript IV reverse transcriptase (ThermoFisher). PCR was carried out using KOD-FX NEO under the following conditions: 120 s at 94°C, followed by 35 cycles of 10s at 98°C, 20s at 60°C, and 60s at 72°C; and 90 s at 72°C, using the primers, P.r_spaca4l_forward and P.r_spaca4l_reverse. Primer sequences were listed in Table S3. The extracted PCR products were sequenced using Big Dye kit 3.1 ver (ThermoFisher) with the Sequencing Service of Center for Gene Research, Nagoya University. The obtained sequence was registered to the DNA Data Bank of Japan (DDBJ, LC722160.1).

### In situ hybridization

Guppy bouncer was cloned as a hybridization probe using RT-PCR. The RT-PCR was performed using KOD-FX NEO (TOYOBO) under the following conditions: 120 s at 94 °C; followed by 35 cycles of 10 s at 98 °C, 20 s at 60 °C, and 60 s at 72 °C; and 90 s at 72 °C, using the primers, P.r_spaca4l_T3_forward and P.r_spaca4l_T7_reverse. The primer sequences are listed in Table S3. The PCR products were used as templates for transcription reactions using T3 RNA polymerase (Biolabs, M0378S) or T7 RNA polymerase (TOYOBO, TRL-201) at 37 °C for 2 h. After transcription, PCR products were digested using the rDNAase kit (QIAGEN, 79254). The RNA strands were then precipitated and eluted with RNAse-free H2O, and the RNA concentration was measured using the Qubit 4 fluorometer (Thermo Fisher Scientific). The paraffin-embedded ovaries were sliced into sections of thickness of 10 µm. Thin sections were mounted on silane-coated glass slides and dried at 37 °C overnight. Then, the sections were deparaffinized in xylene for 10 min; stepwise dehydrated in 100, 75, 50, 25, and 0% ethanol/PBS; pre-fixed in 4% PFA/PBS and washed 3 times with PBS for 5 min; treated with 20 µg/mL ProK (Invitrogen, 0061322)/PBS; post-fixed in 4% PFA/PBS and washed 3 times with PBS; and acetylated in 10 mM triethanolamine with acetic anhydride (20 min, RT). The sections were washed twice with 4X SSC (2 min, RT). The primary wash was performed with 50% formamide/2X SSC (1 h, 65 °C). Slides were pre-hybridized (1 h, 65 °C) with the hybridization buffer (50% formaldehyde, 5X SSC, 200 µg/mL yeast RNA extraction, and 500 units heparin) and hybridized with 100 ng/m each probe/hybridization buffer (16 h, 65 °C). After hybridization, primary wash was performed with 50% formamide/2X SSC (1 h, 65 °C) and secondary wash with 0.1 X SSC (2 h, 65 °C), and the solution was replaced with TTBS; then, the slides were blocked for 30 min in Blocking-One. Antibody reaction was performed with 5000X diluted anti-AP antibody (Sigma-Aldrich 11093274910) (2 h, RT); then, the slides were washed for four times with TTBS (15 min, RT), followed by NTM (10% 1mM pH 9.5 Tris-HCL, 5% 1 mM MgCL, and 2% 5 mM NaCl) (15 min, RT). Colorization was performed using 337.5 µg/mL NBT/ 175 µg/mL BCIP/NTM solutions. Then, the slides were washed with PBS (3 min, RT), fixed in 4% PFA/PBS (10 min, RT), and washed with PBS three times (3 min, RT). After washing, tissues were stained with Nuclear Fast Red (Cosmo Bio, NFS500) and washed with water. After staining, the slides were dehydrated with ethanol, permeabilized with xylene, and embedded in Entellan^TM^ New (Merk).

### Phylogenetic analysis

Amino acid sequences of bouncer, spaca4, and spaca4l from guppy, medaka, and zebrafish were collected from the NCBI protein database (https://www.ncbi.nlm.nih.gov/protein).

Phylogenetic trees were constructed using the neighbor-joining method, with 500 bootstrap replicates using the MEGAX (version 10.1.8) software (https://www.megasoftware.net/) [28, 29]. Accession numbers of the genes used in the phylogenetic analysis are described in Figure 3C.

### Decoy synthesis

Guppy or zebrafish bouncer without a signal peptide and transmembrane region was cloned as a decoy-coding sequence using 2-step RT-PCR. In the first PCR step, ribosome binding sequence was added to the 5′ end of the forward primer, P.r_spaca4l_RBS_forward or D.r_bouncer_RBS_forward, for *in vitro* transcription and translation (Table S3). The sequence for 6 x His was added to the 5′ end of the reverse primer, P.r_spaca4l_His_reverse or D.r_bouncer_His_reverse, for purification of the protein product (Table S3). PCR was performed using KOD-FX NEO under the following conditions: 120 s at 94 °C; followed by 35 cycles of 10 s at 94 °C, 20 s at 60 °C, and 60 s at 72 °C; and 300 s at 72 °C. In the second PCR step, T7 promoter sequence was added to the 5’ end of the forward primer, T7promoter_SD_primer. The sequence for 6 x His was added to the 5’ end of the reverse primer for purification of the protein product. PCR was performed using KOD-FX NEO under the following conditions: 120 s at 94 °C; followed by 35 cycles of 10 s at 94 °C, 20 s at 60 °C, and 60 s at 72 °C; and 300 s at 72 °C.

The *in vitro* transcription and translation were performed using the PureFrex 2.1 kit (GeneFrontier, PF213-0.25) with the PDI set (GeneFrontier, PF006-0.5). The experimental procedure was performed in accordance with the standard protocol [30]. The protein products were purified using the NEBExpress Ni Spin Columns (New England Biolabs, S1427S). The concentration of the purified products was quantified using the Qubit protein assay kit (Thermo Fisher, Q33211). The protein products were diluted to 1 µg/mL in L-15 and stored at -80 °C until use.

### Decoy assay

Female guppies, medaka, and zebrafish were anesthetized in ice water, and the ovaries were removed. Immature eggs were isolated from the ovaries. Male guppies, medaka, and zebrafish were anesthetized with ice water, and the testes were removed and homogenized. Each homogenate was stained with SYTO9 (Invitrogen, 2397781) for 10 min and washed three times with PBS. Washed spermatozoa were co-cultured with 1 µg/mL synthesized bouncer/L-15 for 30 min at RT. Spermatozoa and immature eggs were co-cultured in 1 mL of L-15 for 30 min at RT. Immature eggs were washed three times with PBS and fixed in 4% PFA/PBS for 30 min. Immature oocytes were washed three times with PBS and observed under the M205FA fluorescence stereomicroscope (Leica).

### Generating transgenic zebrafish

The guppy bouncer coding sequence without a stop codon was cloned from the ovary sample and inserted between the *Bgl*II and *EcoR*I sites of the pEGFP-N1 plasmid (Clontech, #6085-1). A nucleotide coding 2A self-cleaving peptides derived from Thosea asigna virus was artificially synthesized and added to the 3’ terminal of the bouncer coding sequence by insertion between the *EcoR*I and *Sal*I sites. The bouncer-2A-GFP coding sequence was amplified using the primers Bouncer_T2A_EGFP_forward and Bouncer_T2A_EGFP_reverse. As a negative control, an intact GFP-coding sequence was amplified using the GFP forward primer and Bouncer_T2A_EGFP_reverse. The zebrafish *gdf9* promoter sequence was amplified using primers D.r_gdf9_promoter_forward and D.r_gdf9_promoter_reverse (Table S3). All PCRs were performed using KOD-FX Neo (TOYOBO). The fragments were inserted within the *EcoR*V site of the *Tol1*-based transgenesis vector pDon122 (Cosmo Bio, CSR-CT-NU-002-1) using the NEBuilder HiFi DNA Assembly System (New England Biolabs). The Tol1-based transgenic construct (5 ng/µL) and Tol1 mRNA transcribed from pHel105 (25 ng/µL; Cosmo Bio, CSR-CT-NU-002- 1) were co-injected into fertilized zebrafish eggs at one- to eight-cell stages. Transgene transcription and protein expression were confirmed by RT-PCR and microscopic detection of GFP.

### Reverse-transcribed PCR

Total RNA was extracted from adult ovaries of transgenic or wild-type zebrafish using the RNeasy Mini kit (QIAGEN, 74134) and reverse-transcribed using SuperScript IV reverse transcriptase (ThermoFisher, 18091050). PCR was performed using KOD-FX NEO under the following conditions: 100 s at 94 °C; followed by 35 cycles of 20 s at 94 °C, 20 s at 60 °C, and 60 s at 72 °C; and 90 s at 72 °C, using the primers, *bouncer*_RT_forward, *bouncer*_RT_reverse, *gfp*-forward, *gfp*-reverse, *actb*-forward, and *actb*-reverse (Table S3).

## Acknowledgements

This study was supported by the Tokai National Higher Education and Research System (THERS) and Interdisciplinary Frontier Next Generation Researcher, and the Japan Science and Technology Agency (JST), and the Japan Society for the Promotion of Science KAKENHI Grant [21H04637 to MK]. Medaka HdrR strain was supplied by NBRP Medaka (https://shigen.nig.ac.jp/medaka/).

## Competing interest

The authors declare no competing interests.

## Author Contributions

J.Y. and A.I. designed the study; J.Y., Y.T., K.U., and A.I. performed the study; M.K., T.S., K.S., and E.H. provided new reagents/analytic tools; J.Y., E.H., and A.I. analyzed data; and J.Y. and A.I. wrote the paper.

## Funding

This work was funded by the Tokai National Higher Education and Research System (THERS) and Interdisciplinary Frontier Next Generation Researcher, and the Japan Science and Technology Agency (JST), and the Japan Society for the Promotion of Science KAKENHI Grant [21H04637 to MK]

## Date availability

All relevant date can be found within the article and its supplemental information.

## Supplemental Information

**Table S1.** Numbers of matured eggs in non-pregnant female.

| Fish ID | Number of mature oocytes |
| --- | --- |
| 1 | 4 |
| 2 | 2 |
| 3 | 2 |
| 4 | 5 |
| 5 | 4 |

**Table S2.** Numbers of littermates after artificial fertilization.

| Fish ID | Number of littermates |  | Total |
| --- | --- | --- | --- |
|  | 1st delivery | 2nd delivery |  |
| 6 | 2 | 7 | 9 |
| 7 | 2 | 4 | 6 |
| 8 | 7 | 0 | 7 |
| 9 | 7 | 0 | 7 |
| 10 | 3 | 0 | 3 |

**Table S3.**
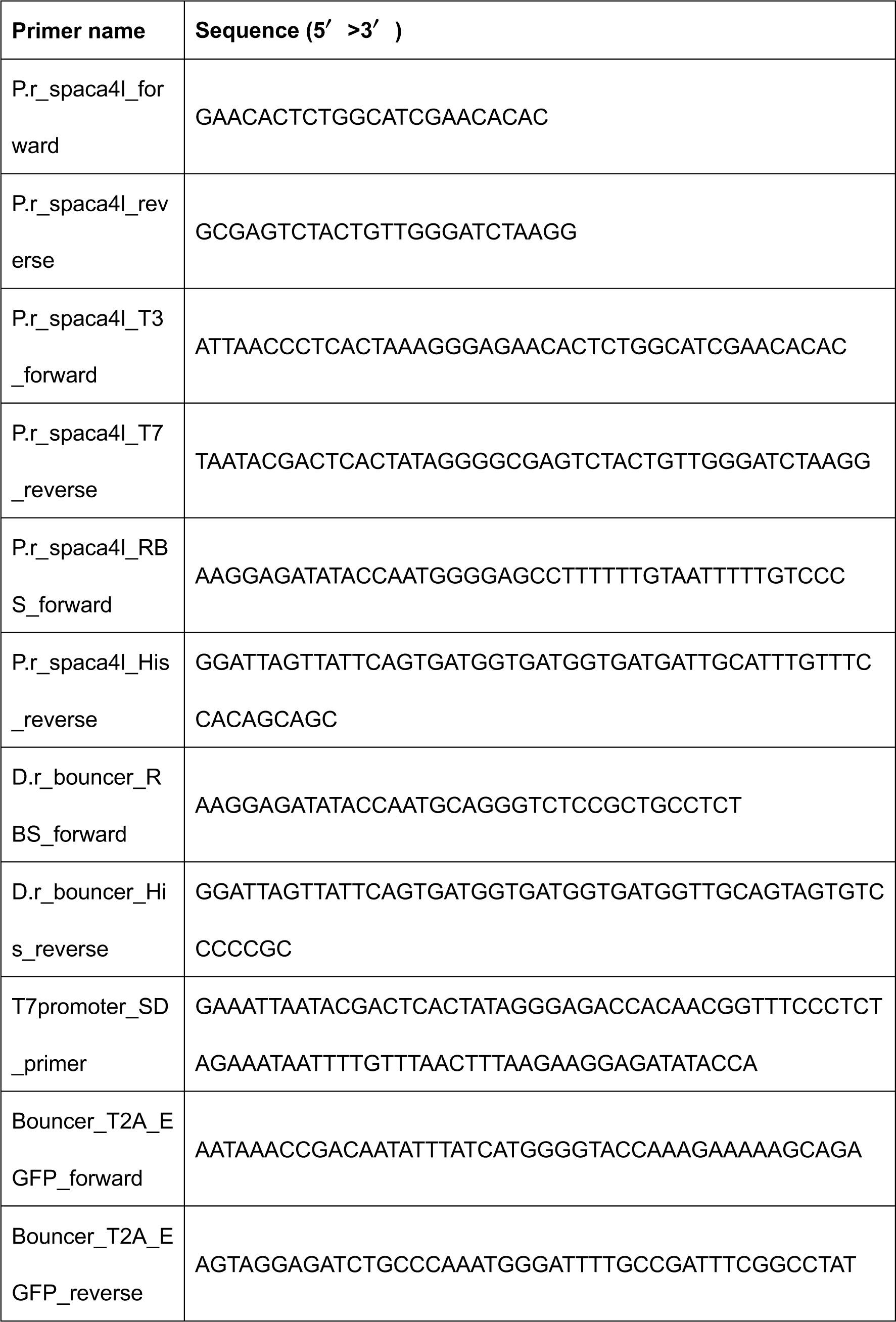
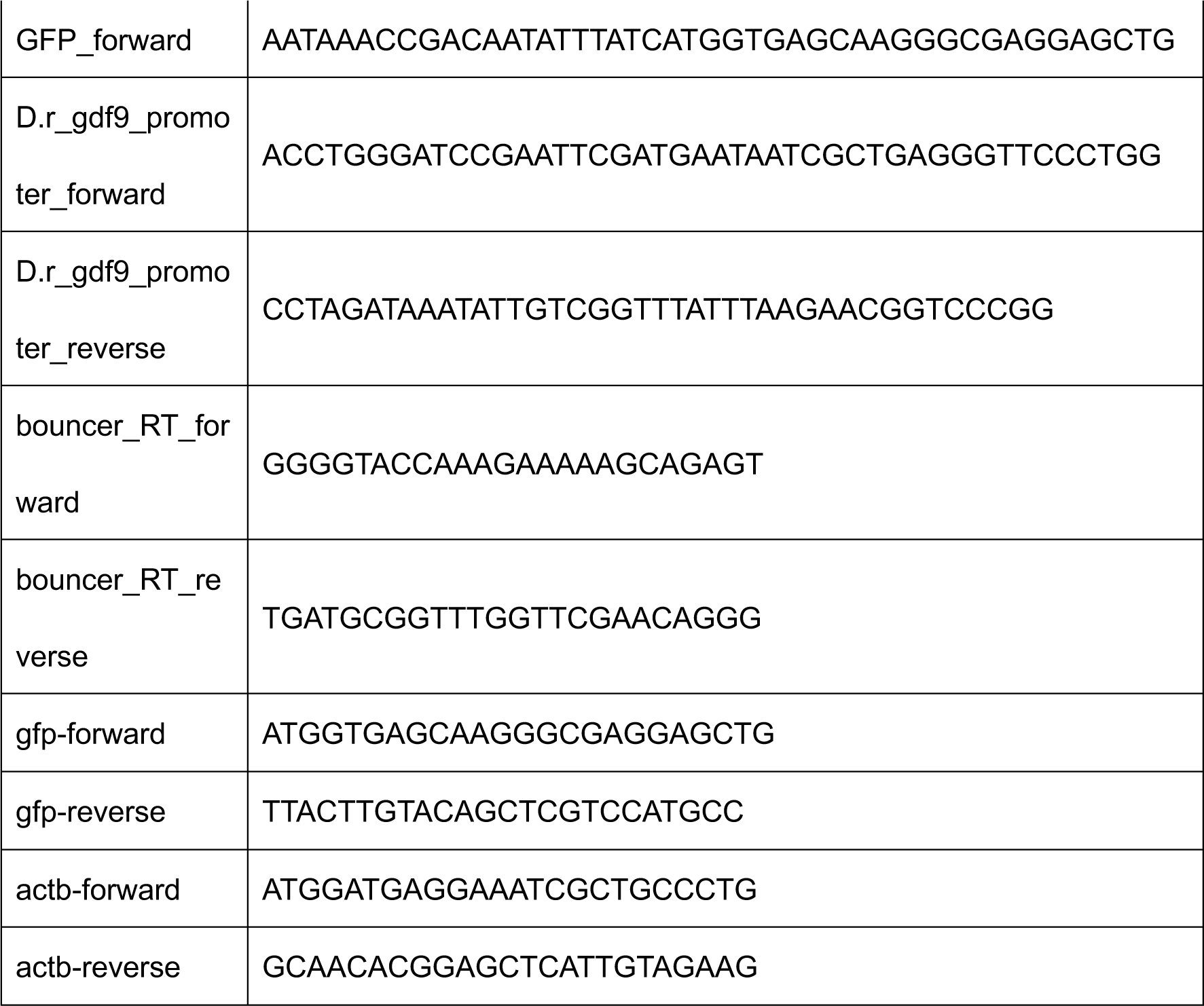
Primers for PCR.

## Supplemental information

**Movie S1.**
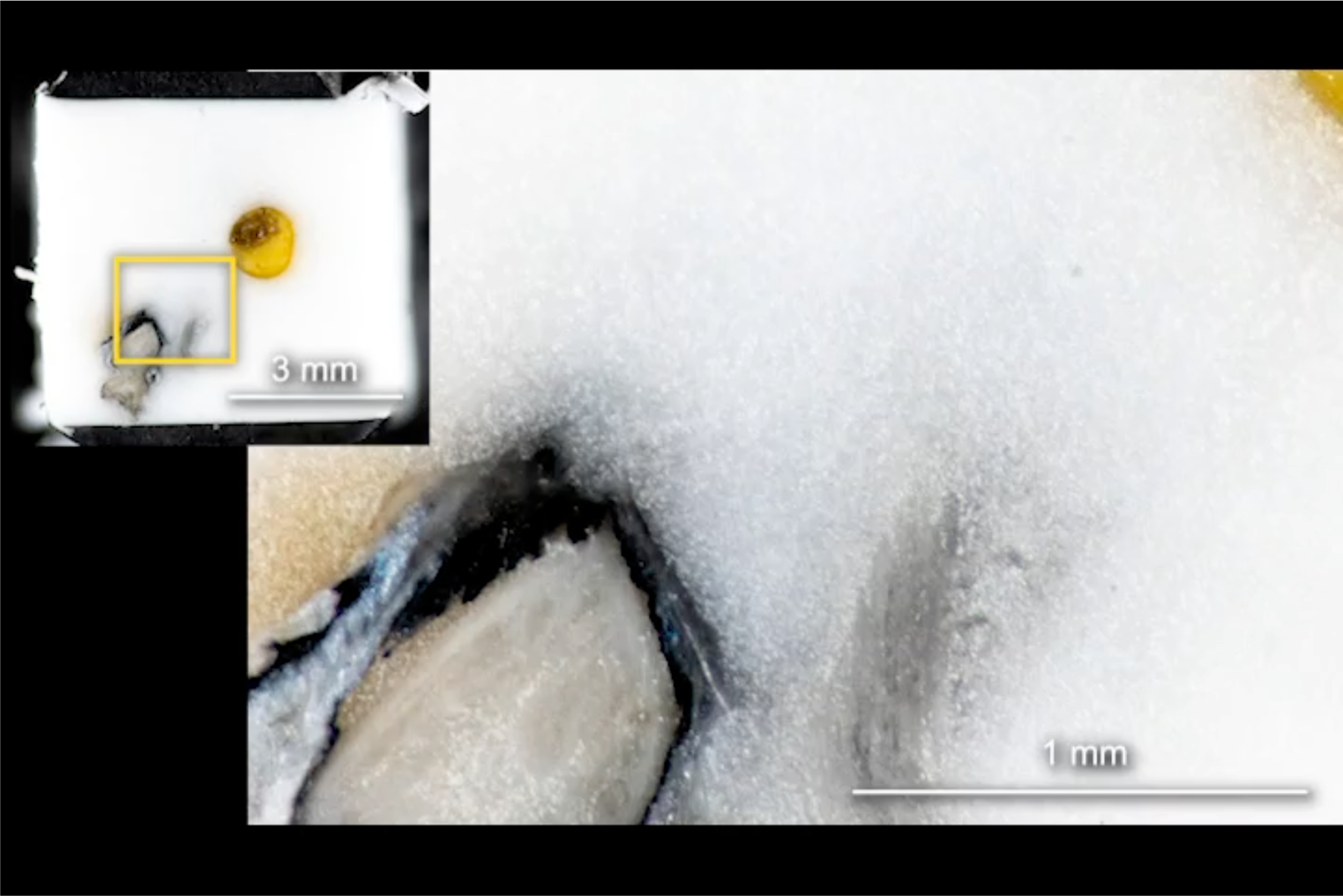
The ducts between the lumen of the ovary and the oocytes were examined by block-face imaging. The ovary of unmated P. reticulata was frozen and sliced at 3 µm thickness. Serial block-face images (1687 images, 1.21 µm/pixel) were obtained using a CoMBI system. Volume-rendered image was generated from the serial block-face images and used to indicate the positions of the planes in Figure1A.

## Notes

### Competing Interest Statement

The authors have declared no competing interest.

## References

1. L. Spallanzani. *Expériences pour Servir A l’Histoire de la Génération des Animaux et des Plantes*. (Barthelemi Chirol, Genève, 1785).

2. P. Talbot, B.D. Shur, D.G. Myles, Cell adhesion and fertilization: steps in oocyte transport, sperm-zona pellucida interactions, and sperm-egg fusion. Biol. Reprod. 68, 1–9 (2003).

3. K. Georgadaki, N. Khoury, D.A. Spandidos, V. Zoumpourlis, The molecular basis of fertilization (Review). Int. J. Mol. Med. 38, 979–986 (2016).

4. P.C. Schroeder, P. Talbot, Ovulation in the animal kingdom: a review with an emphasis on the role of contractile processes. Gamete Res. 11, 191–221 (1985).

5. C. Thibault, D. Szöllösi, M. Gérard, Mammalian oocyte maturation. Reprod. Nutr. Dev. (1980) 27, 865–896 (1987).

6. R.A. Wallace, K. Selman, Cellular and dynamic aspects of oocyte growth in teleosts. Am. Zool. 21, 325–343 (1981).

7. G. Albert. Catalogue of the Fishes in the British Museum. Vol. 6. (Taylor & Francis, London, 1866), p. 353.

8. L. Parenti, A phylogenetic and biogeographic analysis of cyprinodontiform fishes (Teleostei, Atherinomorpha). Bull. Am. Mus. Nat. Hist. 168, 335–557 (1981).

9. J.D. Reynolds, M.D. Gross, M.J. Coombs, Environmental conditions and male morphology determine alternative mating behaviour in Trinidadian guppies. Anim. Behav. 45, 145–152 (1993).

10. M.C. Uribe, H.J. Grier, Insemination, intrafollicular fertilization and development of the fertilization plug during gestation in Heterandria formosa (Poeciliidae). J. Morphol. 279, 970–980 (2018).

11. M.C. Uribe, G. De la Rosa Cruz, A. García Alarcón, J.C. Campuzano Caballero, M.G. Guzmán Bárcenas, Structures associated with oogenesis and embryonic development during intraovarian gestation in viviparous teleosts (Poeciliidae). Fishes 4, 35 (2019).

12. R. Yanagimachi in The Physiology of Reproduction. 2nd ed, E. Knobil, J. Neill, Eds. (Raven Press, New York, 1994), pp. 189–317.

13. W.J. Swanson, V.D. Vacquier, The rapid evolution of reproductive proteins. Nat. Rev. Genet. 3, 137–144 (2002).

14. S. Tardif et al., Zonadhesin is essential for species specificity of sperm adhesion to the egg zona pellucida. J. Biol. Chem. 285, 24863–24870 (2010).

15. E. Bianchi, B. Doe, D. Goulding, G.J. Wright, Juno is the egg Izumo receptor and is essential for mammalian fertilization. Nature 508, 483–487 (2014).

16. S. Herberg, K.R. Gert, A. Schleiffer, A. Pauli, The Ly6/uPAR protein Bouncer is necessary and sufficient for species-specific fertilization. Science 361, 1029–1033 (2018).

17. M.I. Binner et al., The Sperm Protein Spaca6 is Essential for Fertilization in Zebrafish. Front. Cell Dev. Biol. 9, 806982 (2022).

18. K.R.B. Gert et al., Divergent molecular signatures in fish Bouncer proteins define cross-fertilization boundaries. Nat. Commun. 14, 3506 (2023).

19. V.D. Vacquier, G.W. Moy, Isolation of bindin: the protein responsible for adhesion of sperm to sea urchin eggs. Proc. Natl Acad. Sci. U. S. A. 74, 2456–2460 (1977).

20. N. Inoue, M. Ikawa, A. Isotani, M. Okabe, The immunoglobulin superfamily protein Izumo is required for sperm to fuse with eggs. Nature 434, 234–238 (2005).

21. S. Kim, C. Spike, D. Greenstein, Control of oocyte growth and meiotic maturation in Caenorhabditis elegans. Adv. Exp. Med. Biol. 757, 277–320 (2013).

22. J.F. Schindler, W.C. Hamlett, Maternal–embryonic relations in viviparous teleosts. J. Exp. Zool. 266, 378–393 (1993).

23. J.C. Trexler, Resource availability and plasticity in offspring provisioning: embryo nourishment in Sailfin mollies. Ecology 78, 1370–1381 (1997).

24. M. Fleuren, J.L. van Leeuwen, B.J.A. Pollux, Superfetation reduces the negative effects of pregnancy on the fast-start escape performance in live-bearing fish. Proc. Biol. Sci. 286, 20192245 (2019).

25. A. López-Sepulcre, S.P. Gordon, I.G. Paterson, P. Bentzen, D.N. Reznick, Beyond lifetime reproductive success: the posthumous reproductive dynamics of male Trinidadian guppies. Proc. Biol. Sci. 280, 20131116 (2013).

26. Y. Tajika et al., A novel imaging method for correlating 2D light microscopic data and 3D volume data based on block-face imaging. Sci. Rep. 7, 3645 (2017).

27. N. Ishii et al., Correlative microscopy and block-face imaging (CoMBI) method for both paraffin-embedded and frozen specimens. Sci. Rep. 11, 13108 (2021).

28. Felsenstein J. CONFIDENCE LIMITS ON PHYLOGENIES: AN APPROACH USING THE BOOTSTRAP. Evolution. 1985;39(4):783–791. doi:10.1111/j.1558-5646.1985.tb00420.x

29. Saitou N, Nei M. The neighbor-joining method: a new method for reconstructing phylogenetic trees. Mol Biol Evol. 1987;4(4):406–425. doi:10.1093/oxfordjournals.molbev.a040454

30. Y. Shimizu et al., Cell-free translation reconstituted with purified components. Nat. Biotech. 19, 751–755 (2001).

